# Defective insulin clearance plays a primary role in the pathogenesis of chronic kidney disease in mice with null deletion of *Ceacam2* gene

**DOI:** 10.1101/2025.11.23.690002

**Authors:** Sivarajan Kumarasamy, Getachew Belew, Agnes O Portuphy, Raziyeh Abdolahipour, Sobia Zaidi, Haixin Yang, Lucia Russo, Simona S. Ghanem, Amira F. Gohara, Ramiro Malgor, Sonia M. Najjar

## Abstract

Endogenous insulin clearance occurs primarily in hepatocytes and to a lower extent in kidney’s proximal tubule cells (KPTCs). CEACAM1 promotes receptor-mediated insulin uptake to be degraded in hepatocytes in a phosphorylation-dependent manner. Its deletion/inactivation causes hyperinsulinemia-driven insulin resistance, steatohepatitis and liver fibrosis. CEACAM2, the dominant CEACAM protein in murine KPTCs, shares a high homology with CEACAM1. Thus, we examined whether it regulates renal insulin disposal to maintain renal homeostasis. KPTCs derived from *Ceacam2* null mice (*Cc2^-/-^*) exhibited lower receptor-mediated insulin uptake. Combined with the gradual decline in CEACAM1-dependent hepatic insulin clearance, impaired renal insulin clearance contributed to chronic hyperinsulinemia and insulin resistance starting at 10 months of age in *Cc2^-/-^*males. This was followed by proteinuria and reduced glomerular filtration rate in association with glomerulosclerosis and tubulointerstitial damage. Increased collagen deposition in *Cc2^-/-^* kidneys could be mediated in part, by hyperinsulinemia-driven activation of the α5β1 integrin-focal adhesion kinase (FAK) signaling pathways. Together, the data demonstrated that loss of CEACAM2 impaired renal insulin clearance that contributed to hyperinsulinemia and resultant insulin resistance, followed by kidney dysfunction and renal fibrosis. This study provided an in vivo demonstration of the regulation of kidney function by insulin clearance along the liver-kidney axis.

## Introduction

The prevalence of Chronic Kidney Disease (CKD) is on the rise, particularly in older adults.^1^ Association of CKD with metabolic dysfunction has been increasingly recognized.^2–5^ The progression of CKD is characterized by key histological features, including glomerulosclerosis and interstitial tubular damage, which are influenced by major risk factors such as hyperinsulinemia with insulin resistance, inflammation, and oxidative stress. Myofibroblastic transformation of mesangial mesenchymal cells causes cell damage. Damaged cells recruit macrophages that release pro-inflammatory (TNFα and IL6) factors, which in addition to increased production of profibrotic factors (such as TGFβ), lead to extracellular matrix remodeling and collagen deposition in the renal interstitium to cause tubulointerstitial fibrosis.^6^

Dysregulated insulin secretion and clearance contribute to chronic hyperinsulinemia. Insulin clearance occurs mostly in hepatocytes and to a lower extent in kidney proximal tubule cells (KPTCs).^7^ CEACAM1, a plasma membrane glycoprotein with a prominent expression in hepatocytes, undergoes phosphorylation by the insulin receptor tyrosine kinase to promote receptor-mediated insulin uptake to be cleared ^8, 9^ and suppress de novo lipogenesis.^10^ Bolstering this role of CEACAM1, its specific deletion or inactivation in liver caused impairment of insulin clearance followed by chronic hyperinsulinemia.^8, 9, 11^ This drove hepatic insulin resistance and hepatic steatosis together with inflammation (steatohepatitis) and ultimately hepatic fibrosis and liver injury.^12–14^

The most prevalent CEACAM proteins in murine kidney is CEACAM2 that shares high homology with CEACAM1, particularly in the amino acid sequence of their intracellular domains (∼95%), including tyrosine (YTVL) and serine phosphorylation sites.^15–17^ Whereas CEACAM1 is highly conserved among species,^18^ CEACAM2 is only expressed in mice (both murine *Ceacam1* and *Ceacam2* genes are the orthologs of human *CEACAM1* gene).^15^ In contrast to CEACAM1, CEACAM2 is not expressed in the liver, but is the dominant CEACAM protein in the kidney where it is restricted to KPTCs. CEACAM2 is also expressed in crypt epithelial cells of the small intestine, and in several brain nuclei, including the ventromedial hypothalamus.^19, 20^ This is consistent with its specific function in the central regulation of energy balance and brown adipogenesis.^20, 21^ Accordingly, male mice with null deletion of *Ceacam2* gene (*Cc2^-/-^*) exhibited hyperphagia, elevated GLP-1–mediated insulin secretion and brown adipogenesis with increased sympathetic nervous system relay to adipose tissue.^20^ Whereas insulin resistance developed in female *Cc2^-/-^* mice at an early age,^19^ it did not develop in the metabolically active male *Cc2^-/-^* nulls ^21^ until about 9-10 months of age when hepatic insulin clearance became impaired owing to the >60% loss in hepatic CEACAM1 expression.^22^ Moreover, male mice also developed high blood pressure with increased dietary salt relative to their wild-type counterparts.^23^

We herein investigated whether loss of CEACAM2 impairs renal insulin clearance in male mice. Given the potential role of chronic hyperinsulinemia and insulin resistance in promoting cardiovascular disease^24^ and kidney dysfunction,^25^ we examined whether *Cc2^-/-^*null males developed spontaneous glomerular and tubular damage that could lead to proteinuria and renal fibrosis in the absence of any environmental trigger.

## Materials and Methods

### Animals

The generation of global *Ceacam2* null mice (*Cc2^-/-^*) and its propagation on C57BL6/J background has been previously described.^19, 21^ Male mice were kept in a 12-hour light–dark cycle having free access to standard chow. All experiments were approved by the Institutional Animal Care and Use Committee at the University of Toledo and at Ohio University.

### Glomerular filtration rate in conscious mouse

As previouly described,^26^ mice were lightly anesthetized (1.5-2% v/v isoflurane) and a small area of flank fur was shaved off. The next day, mice were anesthetized for <5 min (1.5 - 2% v/v isoflurane), and the NIC-Kidney device (MediBeacon Inc, St. Louis, MO) was attached to the back of the mice using a double-sided adhesive gauze tape. After recording the baseline period for ∼1 min, FITC-Sinistrin (7.5 mg/100g body weight; dissolved in physiological saline solution) was injected into a mouse via retro-orbital route. Each mouse was placed into an individual cage to minimize the risk of probe dislodgement. After a one-hour recording period, the device was carefully removed, and the data was analyzed using NIC-Kidney device partner software (MediBeacon, St. Louis, MO, USA). All mice had *ad libitum* access to food and water except during the one-hour period of measuring glomerular filtration rate (GFR).

### Urinary protein excretion measurement

Mice were housed individually in metabolic cages, and urine samples were collected overnight. Urinary protein excretion was measured as described previously.^27^

### Morphometric analysis

Kidney paraffin sections (3-4 µm thick from 4-5 mice/genptype/age group) were stained with Masson’s trichrome, periodic acid–Schiff (PAS) and H&E for assessment of mesangial expansion and tubulointestitial alterations under light microscopy, as described.^28, 29^ In brief, the severity of glomerulosclerosis was evaluated in at least 30 glomeruli. The extent of damage was assessed by determining both the number of glomeruli showing sclerosis and the sclerotic area within each affected glomerulus. The glomeruli were graded on a scale of 0 to 4: grade 0: normal; grade 1: <25% involvement of the glomerular tuft; grade 2: 25–50% involvement; grade 3: 50–75% involvement; and grade 4: sclerosis occupying >75% of the glomerular tuft. The glomerular damage score was calculated as follows: [(1×N glomeruli with Grade 1) + (2×N glomeruli with grade 2 + (3×N glomeruli with grade 3) + (4×N glomeruli with grade4)] / total number of glomeruli examined. The extent of tubulointerstitial damage-including infiltration, fibrosis, tubular dilatation, and tubular regeneration was assessed by assigning a grade based on the percentage of affected area: grade 0: normal; grade 1: <10%, grade 2: 10–25%; grade 3: 25–50%; grade 4: 50–75%, and grade 5: 75–100%. Sections were examined and scored by two pathologists (AFG and RM independently conducted the analysis).

### Biochemical parameters

Following an overnight fast in cages with Alpha-dri bedding, mice were anesthetized with an intraperitoneal (IP) injection of pentobarbital (1.1mg/kg BW). Retro-orbital venous blood was drawn into heparinized micro-hematocrit capillary tubes (Fisherbrand, Waltham, MA) and plasma was processed and stored at -80°C. Plasma was analyzed for insulin (80-INSMSU-E01 ELISA kit; Alpco, Salem, NH), C-peptide (80-CPTMS-E01 ELISA kit; Alpco), Renin (MAK157-1 KT-Sigma Aldrich, USA), Tumor necrosis factor-alpha (TNFα ELISA Kit, Abcam, Cambridge, MA). Interleukin 6 (IL6 ELISA Kit, Abcam), and Endothelin-1 (ELISA Kit, Abcam).

### Isolation of primary kidney proximal tubule cells

KPTCs were isolated from kidneys of 6-month-old, anesthetized mice (n=5/genotype), as previously described.^30^ Kidneys were decapsulated, horizontally cut, and the medulla was discarded while cortex was finely minced by a razor blade in solution 1 (DMEM-F12, 1mM Heptanoate Acid, 4mM Glycine, pH 7.4). Cortical fragments were digested five times (shaking water bath at 37°C in 100% oxygen for 12 min) in 10ml of Collagenase Solution [(1mg/ml) Collagenase type II (Worthington), 1mg/ml insulin-free BSA, 0.1mg/ml DNAse I (Sigma)], and allowed to settle down by gravity into a 10ml sterile pipette. The first digested solution was discarded and the following digested solution containing tubules were collected in 50 ml falcon tubes and kept in ice. The final combined supernatant was centrifuged at 1000rpm at 4°C for 5 min to collect pellet that was reconstituted in Percoll Solution (Sigma) and ultracentrifuged in Nalgene tubes (Thermo Scientific) at 13000rpm at 4°C for 45 min. Proximal tubular cells aggregating in between the third and fourth layer at ∼1.0-1.3g/ml Percoll density were carefully removed and washed twice in 50ml of Solution 1 (1000rpm, 4°C, 5 min) in 50ml falcon tube. They were combined and plated at equal density onto poly L-lysine coated 6-well-plates at 37°C in a humidified 5% CO_2_ air incubator for 2 days in DMEM/F-12 culture medium (Sigma), containing 1% penicillin-streptomycin, 60nM sodium selenite, 1.1mg/ml sodium bicarbonate, 5 µg/ml human apo-transferrin, 2mM glutamine, 50nM dexamethasone, 5pM of 3,5,3’-triiodothyronine, 25ng/ml Prostaglandin E1, 50nM hydrocortisone, 10ng/ml epidermal growth factor, 5µg/ml insulin, 3.1g/l D-glucose, 2% (vol/vol) FBS, and 20mM Hepes, pH 7.4 (all from Invitrogen and Sigma). Cells were kept in the culture media for a total of 24-48 hrs.

### Ligand binding and internalization

Isolated KPTCs were incubated for 36-48 hours prior to allow 30,000cpm [I^125^]insulin (Human I^125^insulin, Perkin-Elmer) to bind to its receptor overnight at 4°C. Insulin internalization was assessed, as previously described.^8^

### Biotin labeling of surface membrane proteins

Isolated KPTCs were grown in DMEM/F-12 media for 24-48 hrs before switching media to low glucose (2.5mM) for 30 min in order to maintain normal expression of CEACAM2.^20^ These cells were first incubated in the absence or presence of 100nM of insulin at 37°C for 5 min, followed by incubation with biotin (1mg/ml) (Pierce Thermo Fisher, Rockford, IL) on ice for 30 min to label cell-surface proteins. Cells were washed with PBS (Ca^2+^and Mg^2+^-100mg/L each and 15mM glycine) followed by cell-lysis for protein extraction. Protein concentration was measured by BCA method, per manufacturer’s instructions (Pierce). 50µg proteins were immunoprecipitated with streptavidin (Thermo Fisher, Waltham, Mass) overnight at 4°C and the pellet washed three times with 1X PBS, and eluted with 1X Laemmli sample buffer (Biorad) before undergoing SDS-PAGE analysis and immunoblotting with a custom-made anti-CEACAM2 (α-mCC2) antibody (anti-peptide polyclonal antibody raised in rabbit against the KLH-coniugated HPLC purified peptide CNAEIVRFVTGTNKTIKGPVH in CEACAM2 (Bethyl Laboratories, Montgomery, TX; dil. 1:50) and insulin receptor (α-IRα) (N20, Santa Cruz).^22^ Loss of biotin-labelled proteins in the presence of insulin relative to its absence indicated internalization of biotin-labeled glycoproteins.

### Western blot analysis

Proteins from kidney lysates and biotinylated KPTCs were electrophoresed onto a 7% SDS-PAGE and immunoblotted with polyclonal antibodies against phospho-insulin receptor beta (pIRβ) (phospho-Y1361) (Abcam), IRβ (C18C4) (Abcam), FAK (Proteintech), phospho-FAK (Cell signaling technology), type1 Collagen (Invitrogen), custom-made affinity purified polypetide antibodies raised in rabbit against custom-made phospho-CEACAM (α-pCC1/2) (Bethyl)^31^, mouse CEACAM1 (α-mCC1),^32^ and CEACAM2 (α-mCC2). Blots were incubated with horseradish peroxidase-conjugated sheep anti-mouse IgG antibody and donkey anti-rabbit IgG antibody (GE Healthcare Life Sciences, Amersham), followed by enhanced chemiluminescence (ECL, Amersham Pharmacia).

### Semi-quantitative Real Time-PCR (RT-qPCR)

Gene expression was measured by real time qPCR (ABI StepOnePlus Real-Time PCR System, Applied Biosystems, Foster City, CA). Total RNA was extracted from frozen kidneys using Nucleospin RNA isolation kit (Macherey-Nagel, Bethlehem, PA, USA). cDNA templates for RT-qPCR were synthesized using 1μg of total RNA, 5X iScript Reaction Mix and iScript reverse transcriptase (Biorad iScript cDNA Synthesis Kit 1708891). RT-qPCR was performed using Fast SYBR Green master mix (Applied Biosystems). mRNA levels were normalized against ribosomal 18s and expressed as fold-change relative to control group. Gene-specific primer sequences are provided in Table S1.

### Statistical analysis

Quantitative data were presented as mean ± standard error of the mean (SEM). Values were analyzed by one-factor ANOVA with Bonferroni correction or Student’s t test. *P<0.05* was considered statistically significant.

## Results

### Reduced renal insulin signaling and internalization in male *Cc2^-/-^*mice

Insulin release during refeeding (FR) of overnight fasted (F) 6-month-old mice caused phosphorylation of the insulin receptor β subunit (IRβ) in kidney lysates of wild-type (*Cc2^+/+^*) mice relative to their basal fasted state (Figure 1A). This caused CEACAM2 phosphorylation, as normalized to CEACAM2 protein levels. The remarkable decrease of renal CEACAM2 during refeeding is reminiscent of its decreased hypothalamic expression during refeeding, and consistent with CEACAM2 role in suppressing food intake.^19, 20^ In contrast, IRβ phosphorylation was lower in kidney lysates from refed *Cc2^-/-^* null mice (Figure 1A).

**Figure 1:**
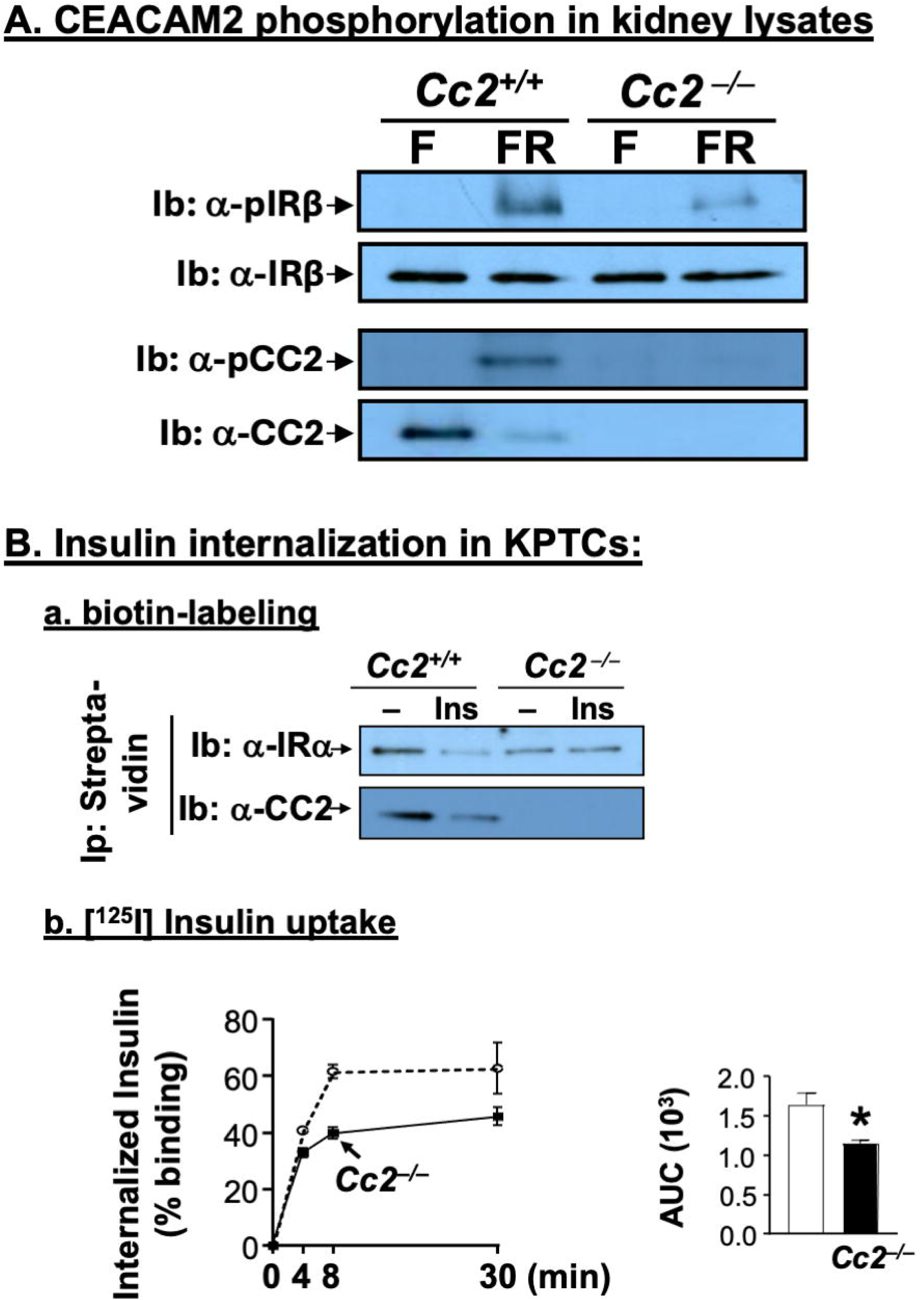
Insulin internalization in KPTCs: **(A)** Mice were overnight fasted (F) and refed (FR) and their kidney lysates subjected to immunoblotting (Ib) analysis with antibodies against phosphorylated IRβ (α-pIRβ) and CEACAM2 (α-pCC2), followed by total α-IRβ, and α-CC2 antibodies as loading controls. **(B)** Primary KPTCs were treated with buffer (–) or insulin (Ins) before cell-surface proteins were labelled with biotin. Proteins were immunoprecipitated (Ip) with α-streptavidin beads prior to analysis by 7% SDS-PAGE and immunoblotting with antibodies against IRα (α-IRα) and CEACAM2 (α-CC2). Lower band density in insulin-treated than buffer-treated wild-type KPTCs refers to reduced cell-surface localization and cellular uptake. **(C)** KPTCs were subjected to 125I-insulin receptor binding at 4°C followed by internalization at 37°C for 0-30 min in triplicate/each time point. The experiment was repeated twice. Area under the curve was assessed and represented in the accompanying graph. Data are expressed as mean ± SEM. *P < 0.05 vs. Cc2+/+ mice.

We then examined whether CEACAM2 loss could impair receptor-mediated insulin internalization in KPTCs, recapitulating the effect of the loss of CEACAM1 in hepatocytes.^8^ As Figure 1B.a shows, immunoblotting (Ib) the biotin-streptavidin immunopellet (Ip) with polyclonal antibodies against IRα and mouse CEACAM2 revealed a decrease in their membrane localization in KPTCs from *Cc2^+/+^*, but not *Cc2^-/-^* mice, when treated with insulin vs buffer alone, indicating insulin-stimulated cellular uptake of insulin receptor and CEACAM2 in wild-type mice. This was supported by lower receptor-mediated [^125^I] insulin internalization in *Cc2^-/-^* relative to *Cc2^+/+^* KPTCs (Figure 1B.b). Thus, receptor-mediated insulin uptake in KPTCs was compromised when CEACAM2 was lost. This translated into reduced insulin clearance in *Cc2^-/-^*relative to wild-type mice, as indicated by the lower C-peptide/insulin molar ratio, beginning at 9-10 months of age, as we have previously reported,^22^ and lasting until 16 months of age (the last age tested) (Figure 2B). The concomitant >60% loss of hepatic *Ceacam1* expression beginning at 10 months (Figure 2B), could contribute to the impairment of insulin clearance at this age. Reduced insulin clearance along the liver-kidney axis in the face of elevated insulin secretion in *Cc2^-/-^* mice ^22^ caused chronic systemic hyperinsulinemia (Figure 3A), as previously observed.^22^

**Figure 2:**
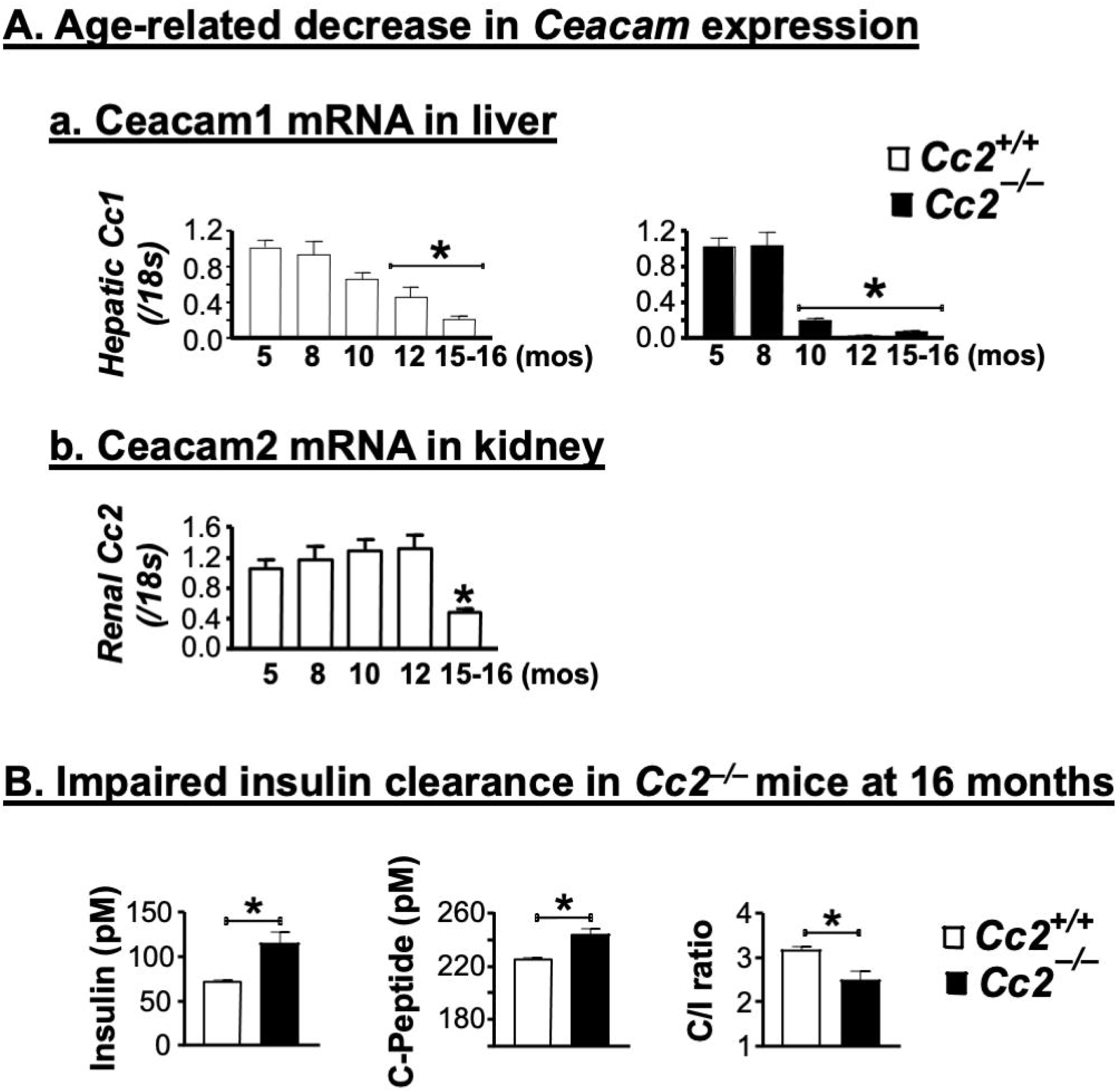
Insulin clearance and action: **(A)** RT-qPCR analysis of mRNA levels of (a) hepatic *Ceacam1* (*Cc1*) and (b) renal *Ceacam2* (*Cc2*) levels normalized to 18s rRNA as internal control. Assay was done in duplicate and data are expressed as fold-change relative to wild-type *Cc2^+/+^* mice (n=4-6/age/genotype). **(B)** Steady-state plasma levels of insulin, C-peptide, and C-peptide/insulin molar ratio (C/I). C/I was calculated as a measure of insulin clearance (n=7-9/age/genotype). Data are expressed as mean ± SEM. *\*P* < 0.05 vs. *Cc2*^+/+^ mice.

**Figure 3:**
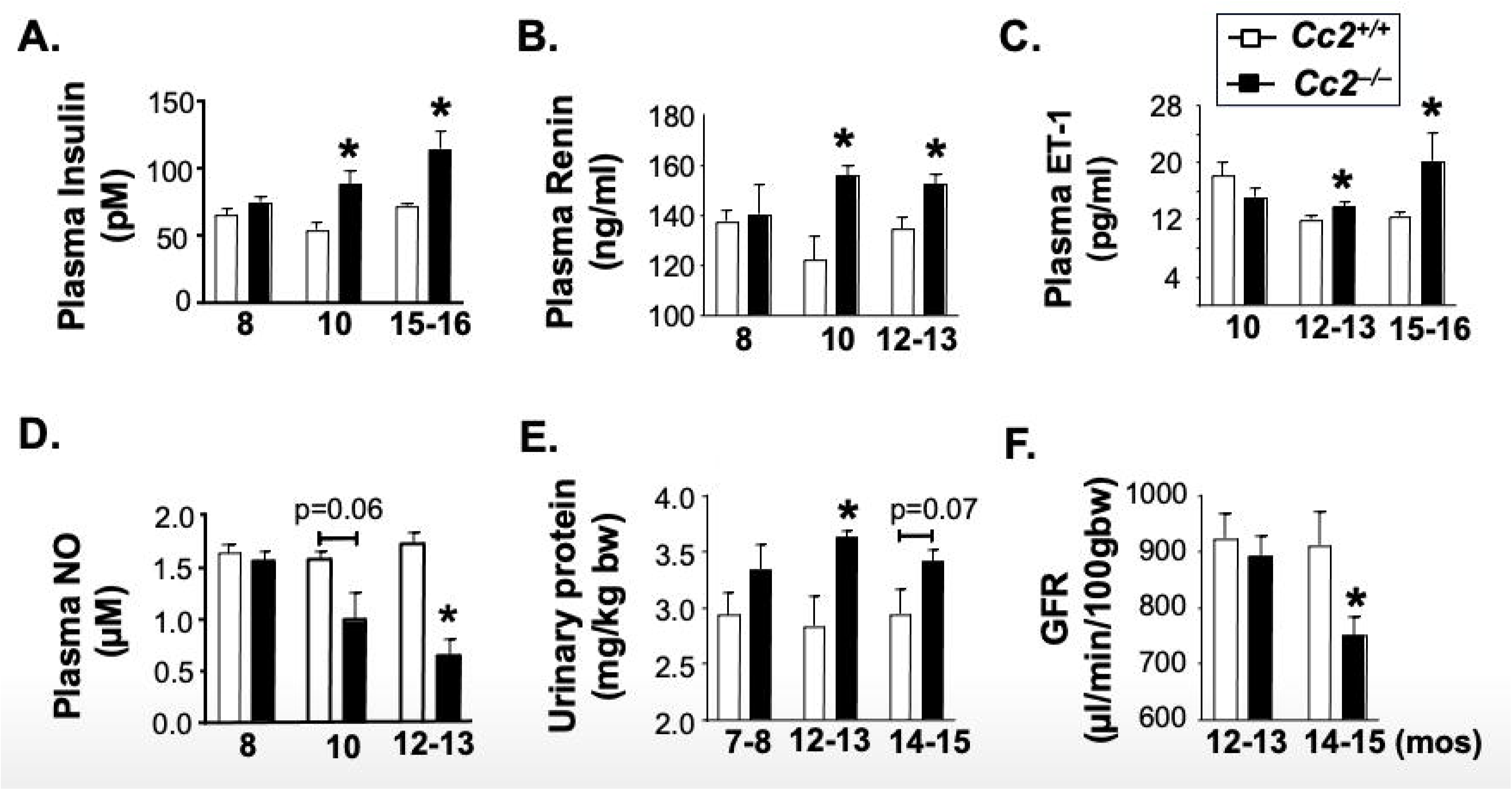
Effect of *Ceacam2* deletion on systemic and renal hemodynamic parameters: **(A-E)** Plasma levels of insulin, renin, ET-1, NO (n=3-9/age/genoptype) and urinary protein excretion (n=5-8/age/genotype) were assayed using commercially available kits. (**F**) glomerular filtration rate (GFR) was assessed by measuring clearance of FITC-Sinistrin injection in conscious mice (n=6-9 mice/genotype) via the fluorescence detector NIC-kidney device. Data are presented as mean ± SEM. \**P* < 0.05 vs. *Cc2*^+/+^ mice.

### Loss of CEACAM2 negatively impacted glomerular filtration rate

Hyperinsulinemia leads to CKD, at least partly by increasing renin levels and activating the renin-angiotensin-aldosterone system (RAAS) in insulin resistance states.^33, 34^ Thus, we evaluated kidney function in *Cc2*^-/-^mice from 4-15 months of age. Concomittantly to hyperinsulinemia (Figure 3A), plasma renin level was higher in *Cc2*^-/-^ nulls than wild-types at 10 months of age (Figure 3B). Consistent with the upregulatory effect of hyperinsulinemia on endothelin-1 (ET-1) production,^35^ mRNA levels of *Et1* and of its *Etar* receptor, which promotes its profibrogenic role, were upregulated by ∼two-to-threefold, while the mRNA level of its *Etbr* anti-fibrogenic receptor was significantly downregulated (by ∼5-fold) in *Cc2*^-/-^ nulls relative to *Cc2^+/+^* mice kidneys (Table S2). Together with the upregulatory effect of hyperinsulinemia on oxidative stress,^24^ plasma ET-1 levels increased (Figure 3C) while nitric oxide (NO) levels reciprocally decreased (Figure 3D) in *Cc2*^-/-^mice starting at 12-13 months of age. This could tip the balance towards vasoconstriction^36, 37^ to contribute to the increase in urinary protein excretion (UPE) at 12-13 months (Figure 3E) preceding a significant decrease in GFR in conscious *Cc2*^-/-^ mice starting at 14-15 months of age (Figure 3F).

### Loss of CEACAM2 drove glomerulosclerosis and tubulointerstitial damage

Histopathological analysis of kidney sections did not show significant differences between *Cc2*^-/-^and *Cc2^+/+^* controls by 11 months of age (not shown). However, histological alterations emerged in mice at >15 months of age. Figure 4 shows representative images of sections from 19-month-old mice stained with H&E (4a and 4d); PAS (4b and 4e) and Masson’s trichrome (4c and 4f). This comprehensive analysis revealed glomerular hypertrophy, hypercellularity, mesangial expansion and basement membrane thickening in *Cc2^-/-^*compared to control *Cc2^+/+^* mice. Combined, these morphological abnormalities yielded a 3-fold increase in the glomerulosclerosis score in *Cc2^-/-^* relative to *Cc2^+/+^* wild-type mice (Figure 4 accompanying graph). Histopathological analysis also revealed inflammation, tubular dilation, tubular regeneration and interstitial fibrosis in *Cc2^-/-^*kidneys compared to *Cc2^+/+^* wild-types at the tubulointerstitial areas (Figure 5). As the accompanying graph shows, the tubulointerstitial damage score was >2-fold higher in *Cc2^-/-^* relative to *Cc2^+/+^*kidneys.

**Figure 4:**
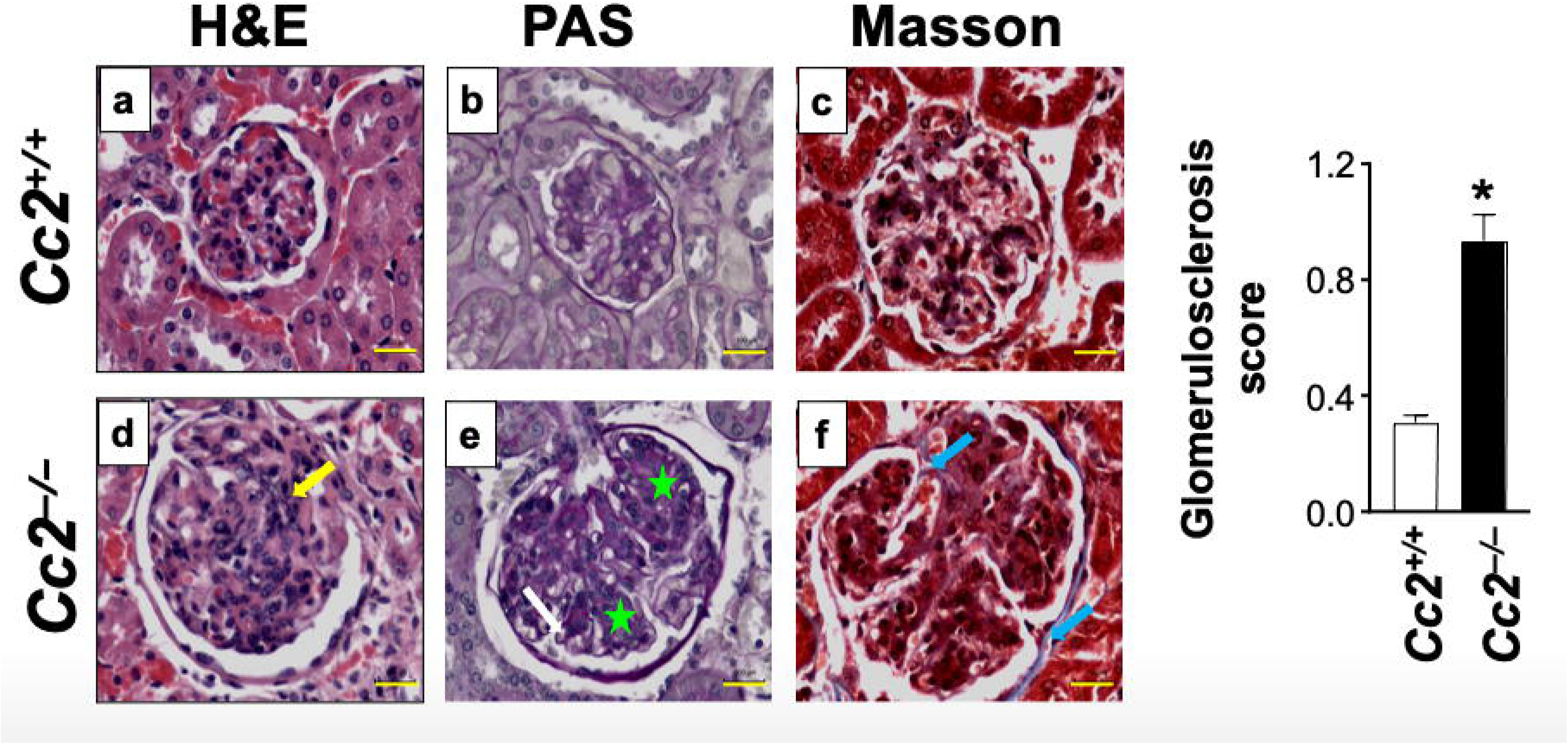
Assessment of glomerular histological alterations: Histopathological analysis revealed glomerular hypertrophy in *Cc2*^-/-^, but not *Cc2^+/+^* kidneys, at 19 months of age (n= 4-5 mice/genotype). H&E staining revealed hypercellularity (yellow arrow; 4d vs 4a), PAS staining showed mesangial expansion (green star; 4e vs 4b) and basement membrane thickening (white arrow; 4e vs 4b) in *Cc2^-/-^* mice. Masson’s trichrome staining revealed mild sclerosis in the glomeruli (blue arrow; 4f vs 4c). Scale bar: 100 μm. Accompanying graph represents the glomerulosclerosis score. Data are expressed as mean ± SEM. *\*P* < 0.05 vs *Cc2^+/+^* mice.

**Figure 5:**
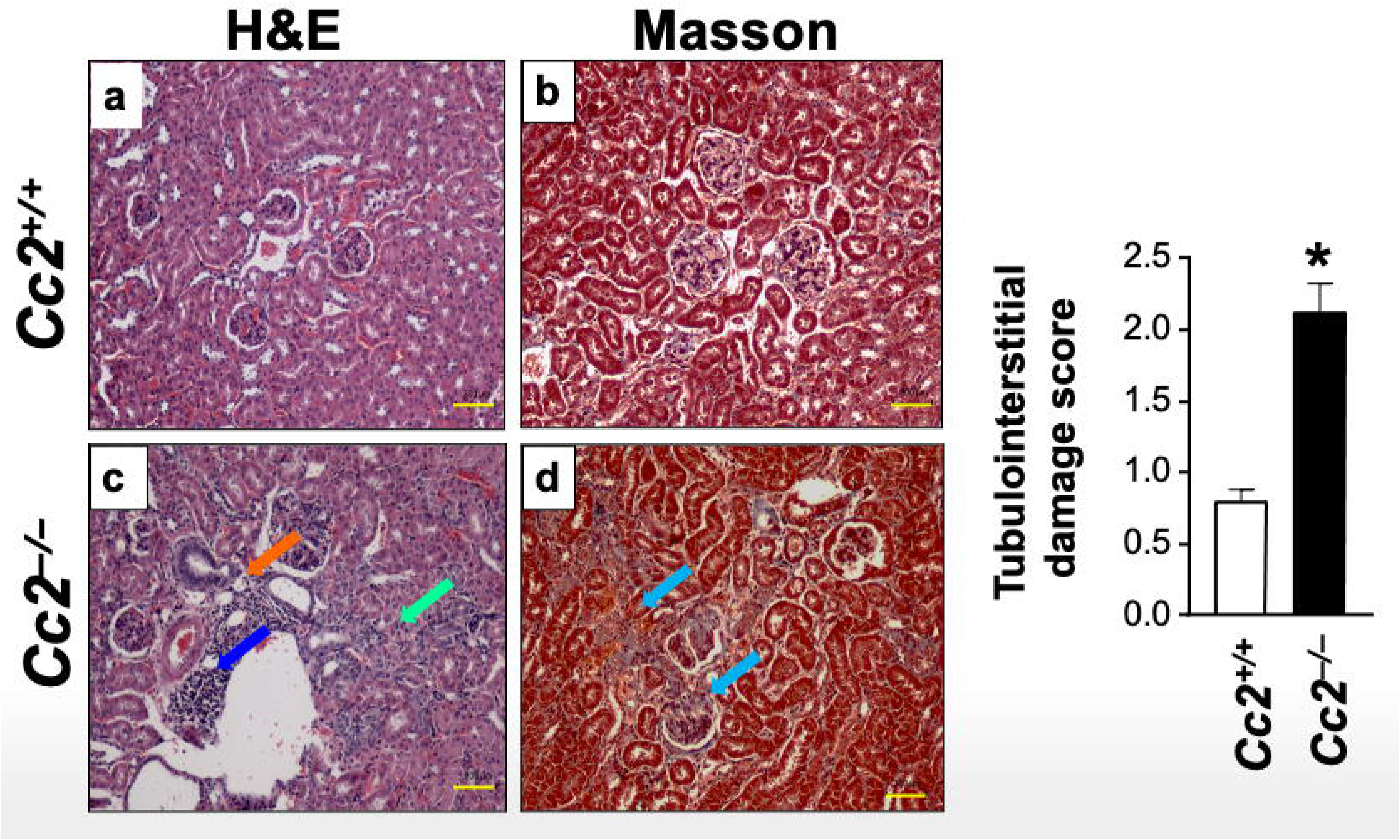
Loss of CEACAM2 promotes tubulointerstitial damage: Histopathological analysis revealed tubular damage in the kidneys of *Cc2*^-/-^, but not *Cc2^+/+^* wild-type mice at 19 months of age (n=4-5 mice/genotype). H&E staining revealed increased tubular dilation (orange arrow; 5c vs 5a), tubular regeneration (green arrow; 5c vs 5a) and accumulation of inflammatory cells (dark blue arrow; 5c vs 5a) in *Cc2*^-/-^ mice. Masson’s trichrome staining demonstrated interstitial tubular fibrosis (blue arrow; 5d vs 5b) in *Cc2^-/-^*, but not *Cc2^+/+^* wild-types. Scale bar: 300 μm. Tubular histology score was represented graphically. Data are expressed as mean ± SEM. *\*P* < 0.05 vs *Cc2^+/+^* mice.

Consistently, the mRNA levels of kidney injury molecule 1 (*Kim1*) was higher in kidney lysates of *Cc2*^-/-^ compared to wild-types beginning at 10 months of age (Table S2) supporting significant proximal interstitial tubular injury. This was accompanied by increased renal mRNA levels of *Tgf*β, followed by elevated mRNA levels of other pro-fibrogenic genes (*Col1a1* and *Col3a1*) and extracellular matrix remodleing (Mmp2, Timp2 and Fibronectin) by ≥12 months of age in *Cc2*^-/-^ than *Cc2^+/+^* kidneys (Table S2), consistent with histological evidence of tubular and glomerular injury in null mice. Together, this demonstrates that *Ceacam2* deficiency exacerbated fibrotic remodeling in the kidney and drove glomerular and interstitial tubular damage linked to proteinuria, reduced GFR, and compensatory glomerular enlargement, eventually leading to kidney insufficiency.

### Loss of CEACAM2 induced a pro-inflammatory microenvironment in kidney

Next, we tested whether *Ceacam2* loss induced a pro-inflammatory state in the kidney given the role of inflammation in the pathogenesis of CKD.^38–40^ *Cc2*^-/-^ mice exhibited an increase in plasma TNFα and IL6 levels starting at 12 months of age compared to *Cc2^+/+^* mice (Figure 6A). Elevated IL6 could activate (phosphorylate) Stat3 (Figure 6B), providing a positive feedback mechanism on the transcription of IL6, TNFα, and other cytokines and chemokines. This includes the early induction (by 8 months) of the expression of pro-fibrogenic gene lipocalin2 (*Lcn2*) (Figure 6C) that significantly contributes to the pathogenesis of CKD in mice and humans. ^41, 42^ As Figure 6C revealed, mRNA levels of other pro-inflammatory markers were induced by 10 months [chemokine ligand 2 and 5 (*Ccl2*, *Ccl5*), interleukin 1β (*Il-1*β), and *Cd68*] followed by *Nlrp3* at 12-13 months of age in *Cc2*^-/-^ compared to *Cc2^+/+^* kidneys. The observed increase in *Ccl2* and *Ccl5* mRNA suggests enhanced recruitment of immune cells to *Cc2*^-/-^ kidneys, as evidenced by elevated expression of *Cd68*, a marker of macrophage infiltration. Notably, the mRNA level of *Nlrp3*, a well-established pro-inflammatory factor that plays a critical role in tubular atrophy in diabetic nephropathy and glomerulonephritis,^43, 44^ was higher in older *Cc2*^-/-^ kidneys. Upregulation of NLRP3 has been implicated in amplifying an inflammatory response through the activation of IL1β, as well as through the TGFβ signaling axis, which exacerbates inflammation and drives fibrotic processes. Together, this indicates that loss of CEACAM2 stimulated the pro-inflammatory milieu in kidneys to contribute to their injury.

**Figure 6.**
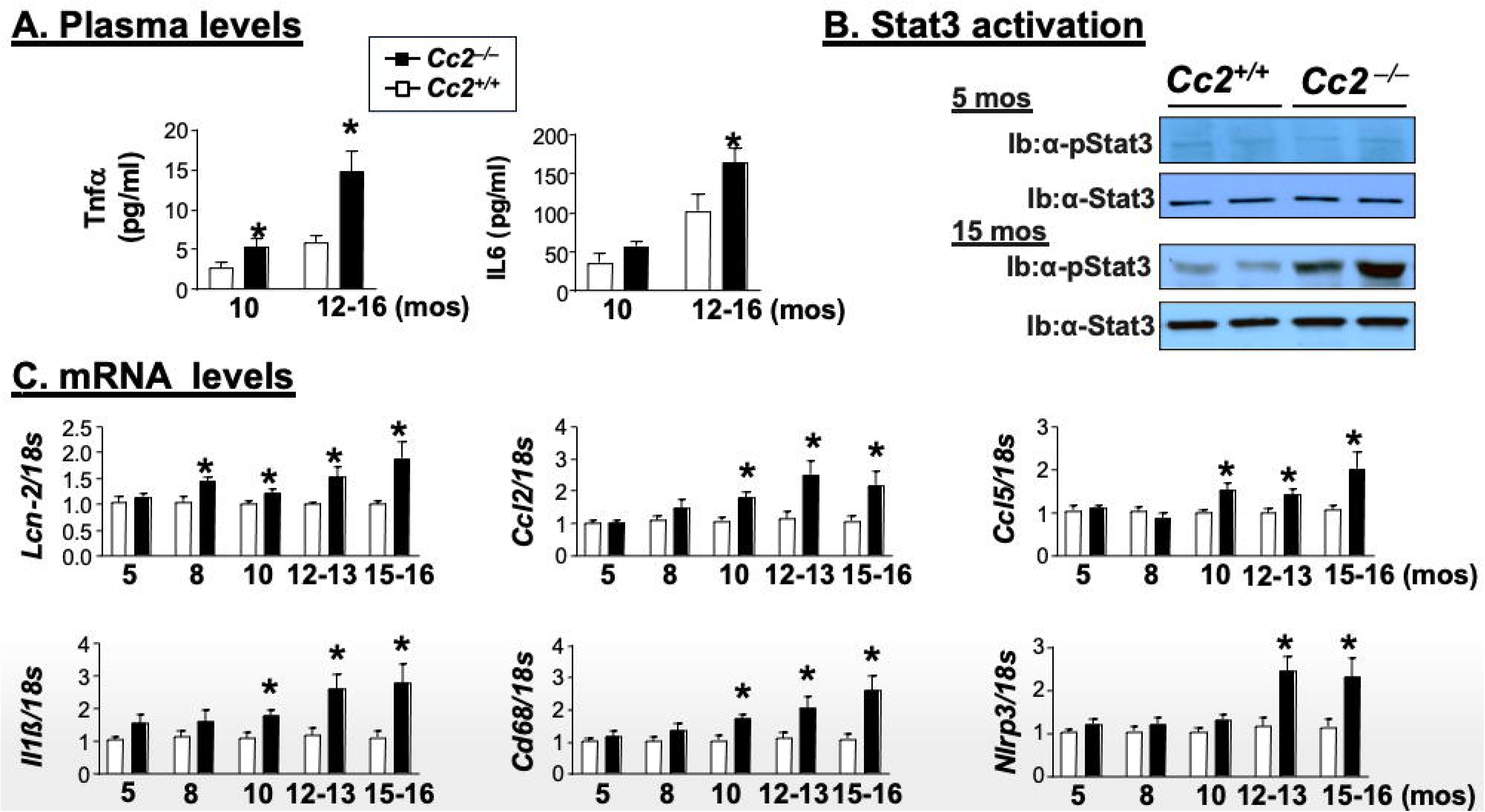
Effect of *Ceacam2* deletion on inflammation: (A) Analysis of plasma levels of TNFα and IL6 was performed in duplicate (n=3-9/age/genotype). Data are expressed as mean ± SEM. *\*P*<0.05 vs *Cc2*^+/+^. (B) Western blot analysis of Stat3 activation (phosphorylation) in kidney lysates from 2 randomly selected mice genotype at 5 and 15 months of age, using antibodies against α-pStat3 and α-Stat33 as loading control. (C) RT-qPCR analysis of mRNA levels of inflammatory markers in the kidneys were done in duplicates (n=3-6/age/genotype), and the expression levels were normalized to 18s rRNA. Gene expression data were expressed as fold-change relative to wild-type *Cc2^+/+^* mice. Data are expressed as mean ± SEM. *\*P* < 0.05 vs. *Cc2^+/+^* mice.

### Loss of CEACAM2 promoted renal fibrosis via activating integrin-FAK signaling pathway

Hyperinsulinemia induces type 1 collagen synthesis in hepatic stellate cells by activating the α5β1 integrin/FAK signaling pathway.^45^ Thus, we tested whether integrin/FAK pathway is activated in *Cc2^-/-^* kidneys. Western blot analysis revealed ∼2-fold increase in β1 integrin protein levels and induced the activation (phosphorylation) of FAK in *Cc2^-/-^* kidney lysates relative to wild-types at 15 but not at 5 months of age (Figure 7). This led to a much higher level of type 1 collagen synthesis in the kidneys of 15-month-old *Cc2^-/-^*, but not wild-type mice.

**Figure 7:**
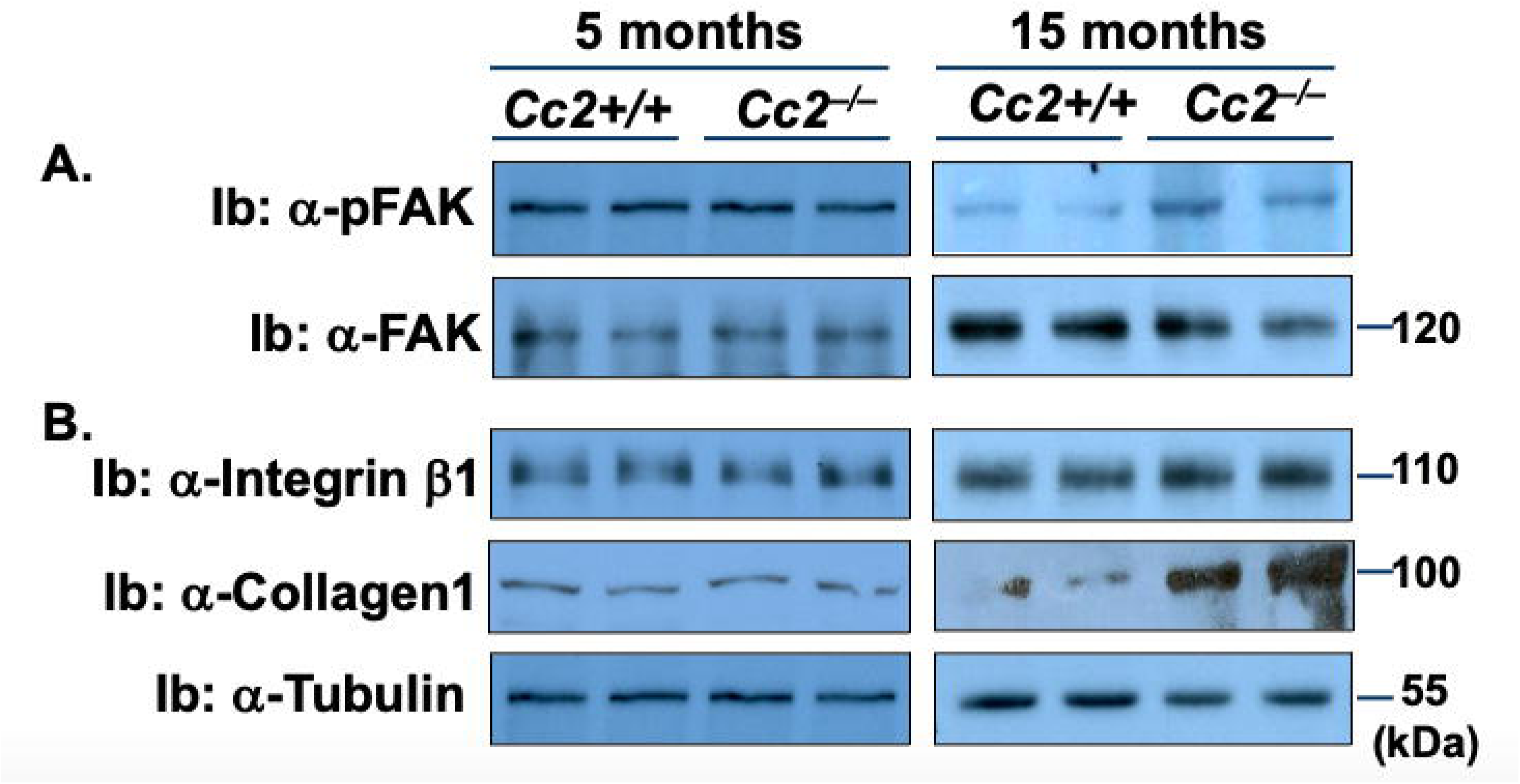
Integrin-dependent FAK signaling in kidney lysates: Western blot analysis of integrin/FAK activation (phosphorylation) in kidney lysates from 2 randomly selected mice/genotype at 5 and 15 months of age. As in the legend to Figure 6, FAK phosphorylation was attained by immunoblotting (Ib) with α-pFAK and normalized against α-FAK antibodies. Protein levels of integrin β1 and type 1 collagen were normalized against α-Tubulin.

## Discussion

Several cross-sectional and clinical research studies pointed to insulin resistance as an independent risk factor for CKD in individuals without diabetes, warranting the identification of underlying mechanisms.^46–48^ The current studies revealed that *Ceacam2* ablation caused glomerulosclerosis and tubulointerstitial damage with progression of kidney dysfunction (proteinuria and low GFR). Renal dysfunction followed the development of hyperinsulinemia and insulin resistance caused by defective insulin clearance in parallel to the loss of CEACAM proteins along the liver-kidney axis. The primary underlying molecular mechanism appears to be defective CEACAM2-dependent receptor-mediated insulin uptake in KPTCs,^7^ providing an in vivo demonstration that impaired renal clearance of endogenous insulin is an independent risk factor of kidney dysfunction. Further supporting the clinical implication of these results is the analysis of v5.nephroseq database showing a progressive decline in the expression of renal CEACAM1 (human ortholog of both murine *Ceacam2* and *Ceacam1*) with increased severity of tubulointerstitial damage in patients with collapsing focal segmental glomerulosclerosis.

Loss of CEACAM2 caused constitutive impairment of renal insulin clearance in KPTCs. Despite increased hyperphagia and GLP-1–mediated insulin secretion, intact insulin clearance in the liver maintained normo-insulinemia, which together with increased energy expenditure, weathered insulin sensitivity in young *Cc2^-/-^* male mice.^22^ With the gradual decrease in energy expenditure as age advanced, fat mass increased with lipolysis-derived free fatty acids that activated PPARα to suppress CEACAM1 transcription in hepatocytes.^49, 50^ Once hepatic CEACAM1 expression was lost by >60% at 9-10 months of age, hepatic insulin clearance became impaired and together with compromised renal CEACAM2-dependent insulin uptake in kidneys, failed to clear excess insulin. This caused chronic hyperinsulinemia with resultant insulin resistance, as manifested by reduced glucose disposal in response to exogenous insulin, random hyperglycemia and blunted insulin signaling in liver and skeletal muscle at this age.^22^ As age advances, the level of insulin degrading enzyme (IDE), a main mechanism of renal insulin clearance, decreases,^51^ likely owing to decreased PPARγ levels.^52^ Combined with the parallel age-dependent decrease in PPARγ-mediated hepatic CEACAM1 expression,^53^ this impaired insulin clearance along the liver-kidney axis to maintain a state of chronic hyperinsulinemia that plays a primary role in CKD pathogenesis in older *Cc2^-/-^*mice.

Whether genetic ablation of *Ceacam1* in hepatocytes could reciprocally cause CKD following impaired insulin clearance in liver is not known, but it is intriguing that receptor-mediated renal insulin clearance increases to compensate for compromised hepatic insulin clearance to limit peripheral hyperinsulinemia in response to high-fat intake.^7^ Because prolonged high-fat intake simultaneously suppresses the expression and activity of IDE,^52^ compensatory increase in receptor-mediated renal insulin clearance is likely mediated by elevated CEACAM2-dependent insulin uptake. While this remains to be investigated, our current findings in *Cc2^-/-^* mice underscore the importance of the bidirectional compensatory mechanism of insulin clearance along the liver-kidney axis in limiting chronic hyperinsulinemia and subsequently, its metabolic and pathological sequelae.

Hyperinsulinemia induces RAAS activation ^54^ and promotes the synthesis of pro-fibrogenic ET-1 production.^35, 55^ It also causes oxidative stress and decreases NO bioavailability.^24^ Consistently, *Cc2^-/-^* mice developed higher plasma ET-1 levels starting at 12-13 months of age. This could derive from increased cell-autonomous production in kidney, as suggested by the increase in renal *Et1* mRNA levels at this age. Together with the induction of the mRNA levels of its pro-fibrogenic *Etar* receptor with the reciprocal reduction of its anti-fibrogenic *Etbr* receptor^55^, this could contribute signifcantly to the fibrogenic processes in *Cc2^-/-^*mice kidneys. Moreover, elevated plasma ET-1 with the reciprocal decrease in NO levels could cause vasoconstriction, as evidenced by the increase in proteinuria and compromised GFR.^55^ Thus, the low GFR reported in *Cc2^-/-^* mice could at least in part, be attributed to the rise in ET-1 production, likely driven by hyperinsulinemia.

Hyperinsulnemia also induces type 1 collagen production by activating α5β1 integrin/FAK pathways in stellate cells to cause hepatic fibrosis.^45^ Consistently, type 1 collagen systhesis was elevated in the kidneys of 15-month-old, but not 5-month-old *Cc2^-/-^*mice, in parallel to activated integrin/FAK pathway. Increased collagen production in turn, could lead to mesangial expansion, a hallmark of glomerular injury.^56^ These observations are supported by activation of TGFβ-mediated fibrotic signaling by α5β1 integrin in fibroblasts.^57, 58^ Collagen synthesis is also regulated by Stat3 activation to contribute to glomerular nephropathy.^59^ The rise of plasma IL6 levels starting at 12 months of age could activate Stat3 in the kidneys of 15-month-old *Cc2^-/-^*mice to contribute to collagen production and cell proliferation, leading to mesangial expansion and thickening of the glomerular basement membrane, as supported by elevated levels of *Kim1* and *Lcn2* mRNA levels.^60, 61^ These findings are consistent with the contribution of hyperinsulinemia in mesangial proliferation, tubulointerstitial remodeling, and extracellular matrix deposition.^25, 62, 63^ Furthermore, hyperinsulinemia could contribute to the development of proteinuria and the progression of CKD via increased TGFβ production in the mesangium and promoting interstitial tubular fibrosis.^63^

In addition to increased IL6 cytokine production, plasma TNFα levels were also elevated in *Cc2^-/-^* mice starting at 10 months of age. Increased plasma TNFα promotes renal vasoconstriction by activating the RAAS pathway,^64^ causing oxidative stress,^65^ and inducing ET-1 synthesis,^66^ thereby decreasing renal blood flow and ultimately reducing GFR. Based on increased coupling of Shc/NF-kB to growth factor receptors in the absence of Shc sequestration by CEACAM1,^67, 68^ increased production of TNFα and IL6 levels in *Cc2^-/-^*mice could predictably result from increased coupling of Shc/NF-kB pathways to the insulin receptor in the absence of CEACAM2.

In addition to this autocrine increase in adipokine production, TNFα and IL6 could also derive from the pro-inflammatory reaction to increased fat mass starting at 9-10 months of age.^22^

Given that the liver and kidney are the two principal sites of insulin clearance, the current findings indicated that reduced insulin clearance resulting from loss of CEACAM proteins along the liver-kidney axis could lead to kidney dysfunction and CKD progression driven by hyperinsulinemia with its resultant insulin resistance and associated pro-inflammation and oxidative stress.

## Acknowledgment

The authors wish to thank Dr. Elsaid Salaheldeen for his help in maintaining the mouse colony, urine collection and for helpful discussions. We also thank Dr. Christian K Raphael and Dr. Harrison T Muturi for his assistance in insulin uptake in kidney proximal tubule cells, Dr. Marziyeh S Jahromi for her assistance in mRNA expression studies, and Dr. Basel G Aldroubi for his assistance in histology analysis.

## Credit authorship Contribution statement

S.K., G.D.B., A.O.P., R.A., S.Z., H.Y., L.R., and S.S.G., researched data. S.K., planned and organized experiments, collected and analyzed data. A.F.M., and R.M., performed histological analysis. S.K., R.M., and S.M.N., discussed data and edited the manuscript. S.K., and S.M.N. conceived and oversaw the work, including its study design and data analysis, leading scientific discussions and drafting/reviewing/editing the manuscript.

## Nonstandard abbreviations used

CEACAM1: Carcinoembryonic antigen-related cell adhesion molecule 1
*Ceacam1*: murine gene encoding CEACAM1 protein
*Ceacam2*: murine gene encoding CEACAM2 protein
*Cc2^-/-^*: global *Ceacam2* null mouse
*Cc2^+/+^*: wild-type littermates
KPTCs: kidney proximal tubule cells.

