## Supplemental File for "Defective insulin clearance plays a primary role in the pathogenesis of chronic kidney disease in mice with null deletion of *Ceacam2* gene"

Nonstandard abbreviations used. CEACAM1, Carcinoembryonic antigen-related cell adhesion molecule 1; *Ceacam1*, murine gene encoding CEACAM1 protein; *Ceacam2*, murine gene encoding CEACAM2 protein; *Cc2*$^{-/-}$, global *Ceacam2* null mouse; *Cc2*$^{+/+}$, wild-type littermates; KPTCs, kidney proximal tubule cells.

**Table S1. Sequences of gene-specific primers used in RT-qPCR analysis**

| Gene | Forward Primer (5' → 3') | Reverse Primer (5' → 3') |
|---|---|---|
| *18s* | TTCGAACGTCTGCCCTATCAA | ATGGTAGGCACGGCGACT |
| *Ccl2/Mcp1* | GCCCCACTCACCTGCTGCTACT | CCTGCTGCTGGTGATCCTCTTGT |
| *Ccl5* | GCCCTCACCATCATCCTCACT | GGCGGTTCCTTCGAGTGACA |
| *Cd68* | CCTCGCCCTAGTCCAAGGTC | CGATTCGGATTTGAATTTGGGCT |
| *Col1a1* | TAGGCCATTGTGTATGCAGC | ACATGTTCAGCTTTGTGGACC |
| *Col3a1* | CATACCTGGTACCGGTGGTC | CACCGACTTCACCCTTTGGA |
| *Et1* | GGTGGAAGGAAGGAAACTAC | CAAGAAGAGGCAGAAAGGCA |
| *Etar* | TCACCGTCTTGAACCTCTGTGC | GATGGAGACGATTTCAATGGCGG |
| *Etbr* | CAGTCTTCTGCCTGGTCCTC | GGACTGCTTTTCCTCAAACG |
| *Fibronectin* | ACGGTGTCAACTACAAGATCG | GTCTTCCCATCGTCATAGCAC |
| *Il1β* | TCGCTCAGGGTCACAAGAAAC | CATCAGAGGCAAGGAGGAAAAC |
| *Kim1* | TGGTTGCCTTCCGTGTCTCT | TCAGCTCGGGAATGCACAA |
| *Lcn2* | CACCACGGACTACAACCAGTTCGC | TCAGTTGTCAATGCATTGGTCGGTG |
| *MMP2* | CAACGGTCGGGAATACAGCAG | CCAGGAAAGTGAAGGGGAAGA |
| *Nlrp3* | ATTACCCGCCCGAGAAAGG | TCGCAGCAAAGATCCACACAG |
| *Timp2* | GCCAAAGCAGTGAGCGAGAAG | GGGGAGGAGATGTAGCAAGGG |
| *Tgf-β* | GTGGAAATCAACGGGATCAG | ACTTCCAACCCAGGTCCTTC |

**Table S2: RT-qPCR analysis of mRNA levels of genes involved in kidney injury and fibrosis**

| Gene | 8 months | | 10 months | | 12-13 months | | 15-16 months | |
|---|---|---|---|---|---|---|---|---|
| | $Cc2^{+/+}$ | $Cc2^{-/-}$ | $Cc2^{+/+}$ | $Cc2^{-/-}$ | $Cc2^{+/+}$ | $Cc2^{-/-}$ | $Cc2^{+/+}$ | $Cc2^{-/-}$ |
| ***Et1*** | ND | ND | 1.09 ± 0.16 | 1.35 ± 0.18 | 1.14 ± 0.22 | 3.11 ± 0.48[a] | 1.17 ± 0.19 | 2.53 ± 0.29[a] |
| ***Etar*** | ND | ND | 1.07 ± 0.14 | 1.20 ± 0.14 | 1.04 ± 0.10 | 1.57 ± 0.16[a] | 1.05 ± 0.10 | 2.63 ± 0.60[a] |
| ***Etbr*** | ND | ND | 1.03 ± 0.09 | 1.24 ± 0.15 | 1.07 ± 0.16 | 0.66 ± 0.06[a] | 1.05 ± 0.11 | 0.20 ± 0.04[a] |
| ***Kim1*** | 1.04 ± 0.10 | 1.33 ± 0.13 | 1.09 ± 0.13 | 1.74 ± 0.13[a] | 1.12 ± 0.15 | 2.25 ± 0.41[a] | 1.09 ± 0.15 | 2.10 ± 0.37[a] |
| ***Tgf-β*** | 1.05 ± 0.12 | 1.07 ± 0.06 | 1.01 ± 0.04 | 1.35 ± 0.08[a] | 1.01 ± 0.04 | 1.38 ± 0.14[a] | 1.04 ± 0.08 | 1.77 ± 0.29[a] |
| ***Col1a1*** | 1.06 ± 0.13 | 0.88 ± 0.14 | 1.04 ± 0.09 | 1.09 ± 0.07 | 1.05 ± 0.11 | 2.18 ± 0.26[a] | 1.16 ± 0.22 | 2.07 ± 0.32[a] |
| ***Col3a1*** | 1.03 ± 0.09 | 0.82 ± 0.03 | 1.03 ± 0.10 | 0.82 ± 0.10 | 1.03 ± 0.09 | 0.80 ± 0.08 | 1.02 ± 0.06 | 1.82 ± 0.18[a] |
| ***Mmp2*** | ND | ND | 1.00 ± 0.04 | 0.85 ± 0.07 | 1.03 ± 0.08 | 1.62 ± 0.11[a] | 1.02 ± 0.07 | 1.58 ± 0.17[a] |
| ***Timp2*** | ND | ND | 1.09 ± 0.17 | 1.00 ± 0.06 | 1/07 ± 0.05 | 0.89 ± 0.06 | 1.06 ± 0.11 | 1.70 ± 0.10[a] |
| ***Fibronectin*** | ND | ND | 1.04 ± 0.10 | 1.02 ± 0.06 | 1.02 ± 0.06 | 0.81 ± 0.07 | 1.05 ± 0.10 | 2.42 ± 0.34[a] |

RT-qPCR analysis was performed in kidneys in duplicate (n=3-6/age/genotype), and the expression levels were normalized to 18s rRNA. Gene expression data were expressed as fold-change relative to wild-type $Cc2^{+/+}$ mice. Data are expressed as mean ± SEM. *$P < 0.05$ vs. $Cc2^{+/+}$ mice.
